# Nanobody-Functionalized AAV achieves Promoter-independent Neuronal Targeting in the CNS å

**DOI:** 10.64898/2026.09.18.752817

**Authors:** Ajay Extross, Ezequiel Marron Fernandez de Velasco, Yungui He, Kevin Wickman, Daniel Schmidt

## Abstract

Adeno-associated virus (AAV) vectors are widely used for gene delivery to the central nervous system, but natural capsid tropism is broad and cell-type restriction is typically imposed transcriptionally using promoters and enhancers that consume packaging capacity and often drive weak expression. Here, we enhance neuronal targeting directly into the AAV-DJ capsid by ablating its endogenous heparan sulfate proteoglycan (HSPG) affinity and genetically displaying a nanobody against the Group 1 metabotropic glutamate receptor mGluR5 within the VP1 subunit at a permissive capsid loop (T456), generating AAV-m5. Western blot confirmed incorporation of the nanobody-VP1 fusion into assembled capsids. In primary hippocampal neuron cultures, ablating HSPG binding abolished infectivity and nanobody display rescued transduction while restricting GFP expression almost exclusively to mGluR5-positive neurons. Heparin competition assays showed that, unlike wild-type AAV-DJ, AAV-m5 transduction was unaffected by exogenous heparin, confirming that entry occurs independently of HSPG binding. Packaged with a strong constitutive promoter (CAG), AAV-m5 achieved neuron-restricted expression comparable to or exceeding that of wild-type AAV-DJ driven by a neuron-specific promoter (hSyn) and produced negligible expression under an astrocyte-specific promoter (GFAP) despite promoter activity in glia, demonstrating that capsid-level targeting can substitute, or complement, transcriptional restriction. Following stereotactic injection into the mouse hippocampus, AAV-m5 achieved an eight-fold higher proportion of transduced neurons than wild-type AAV-DJ at equivalent titers. Delivery to Grm5-null hippocampus reduced both the intensity and the spatial extent of transduction, confirming that the broad hippocampal transduction achieved by AAV-m5 is mGluR5-dependent. Together, these results establish nanobody-functionalized AAV-DJ as a modular, single-component platform for precision CNS gene delivery that circumvents the packaging and expression trade-offs of promoter-based cell-type restriction, with potential for retargeting to additional CNS cell types and disease-relevant receptors.

## Introduction

Adeno-associated virus (AAV) has become a widely used vector for gene delivery in both research and the clinic, owing to its non-integrating genome, low immunogenicity, and well-characterized biology ^1, 2^. A central constraint of the platform is its limited packaging capacity of approximately 4.7 kb, which bounds the size of any genetic payload it can carry ^1^. Naturally occurring serotypes display broad and overlapping tissue tropism that is dictated largely by interactions between the viral capsid and cell-surface glycans: AAV9, for example, engages terminal N-linked galactose ^3, 4^, whereas the engineered shuffled variant AAV-DJ retains high affinity for heparan sulfate proteoglycans inherited from its AAV2 lineage ^5^. Productive transduction by most serotypes further depends on the universal proteinaceous receptor AAVR (KIAA0319L), which couples initial glycan attachment to cellular entry ^6^.

In neuroscience, AAVs are indispensable for delivering genetic payloads to defined cell populations to dissect and manipulate circuits ^7^. Yet the very breadth of natural tropism that makes these vectors versatile also drives indiscriminate transduction of neurons, astrocytes, and other glia when injected into the brain, confounding experiments and posing risks for therapeutic applications ^7, 8^. The prevailing strategy for restricting expression to a target cell type relies on cell-type-specific promoters and enhancers, such as the forebrain-derived GABAergic interneuron-targeting mDlx enhancer, medium spiny neuron enhancer, and others ^9–12^. This approach carries two persistent drawbacks. First, bulky regulatory elements in some cases consume scarce packaging capacity, forcing a trade-off against the therapeutic or experimental cargo ^13, 14^. Second, cell-type-specific regulatory elements typically drive weaker expression than constitutive promoters such as CAG or CMV, often yielding suboptimal transgene levels ^14, 15^.

To overcome these limits, a complementary strategy seeks to redirect tropism at the level of the capsid itself, so that physical targeting rather than transcriptional control confers specificity. Directed evolution of capsid libraries by error-prone PCR and DNA family shuffling, coupled to *in vivo* selection, has produced variants with markedly altered tropism, including the central nervous system-penetrant AAV-PHP.B and AAV-PHP.eB and the retrograde-access variant AAV2-retro ^5, 16–19^. Rational peptide display, in which short targeting peptides are inserted into surface-exposed capsid loops such as the variable region VIII near residue 588, offers a more directed route to novel binding ^20, 21^. An increasingly powerful alternative grafts defined targeting moieties onto the capsid, evolving from early bispecific-antibody adaptors ^22^ to genetically displayed high-affinity ligands and designed ankyrin repeat proteins (DARPins) that redirect AAV to chosen surface receptors, including in neurons ^23–25^. Among these moieties, nanobodies (single-domain VHH antibodies) are especially attractive because their small size, high stability, and ease of recombinant production allow direct genetic insertion into the capsid ^26^. Genetically encoded nanobodies have been used to retarget AAV to membrane antigens ^27^, and recent work mapping permissive capsid insertion hotspots has shown that a nanobody against fibroblast activation protein (FAP) can confer precise, modular retargeting to FAP-expressing tissues ^28, 29^.

Building on this modular nanobody-display technology, we sought to apply it to a target of high clinical relevance in the central nervous system. Here we target the Group 1 metabotropic glutamate receptor mGluR5, a G protein-coupled receptor implicated in a range of neurodegenerative and psychiatric disorders ^30–32^. By genetically ablating the endogenous heparin affinity of the AAV-DJ capsid and incorporating an mGluR5-specific nanobody at a permissive insertion site, we engineered a vector that achieves high-fidelity neuronal targeting driven by physical tropism rather than by restrictive promoters.

## Results

### Engineering and Structural Validation of Nanobody-Functionalized AAV-DJ

To redirect the tropism of engineered AAV vectors, we sought to ablate endogenous, widespread heparan sulfate proteoglycan (HSPG) binding and replace it with precise, receptor-mediated interactions. The proposed infection mechanism, utilizing a tethered nanobody (Nb) to bypass traditional attachment factors, is depicted in **Figure 1A**. To generate an HSPG-null scaffold, we targeted the heparin-binding domain (HBD) of the AAV-DJ capsid by introducing R587A/R590A substitutions, which have been shown to substantially reduce heparin affinity ^33^. We then genetically incorporated the Nb43 nanobody, which targets the neuronal receptor mGluR5 ^34^, into the VP1 subunit of AAV-DJ at position T456. To balance conformational flexibility with appropriate spatial presentation of the displayed Nb, two linker configurations were evaluated: a short symmetrical (GGGGS)₁ linker and a longer asymmetrical (GGGGS)₅/(GGGGS)₁ arrangement (**Figure 1B**). Successful capsid assembly and Nb incorporation were confirmed by Western blot analysis of iodixanol-purified AAV particles. All three capsid proteins (VP1, VP2, and VP3) were detected; notably, AAV-m5 variants displayed an upward shift of ∼15 kDa in the VP1 band relative to wild-type AAV-DJ (**Figure 1C**), consistent with the predicted mass of the Nb–VP1 fusion (∼96 kDa).

**Figure 1.**
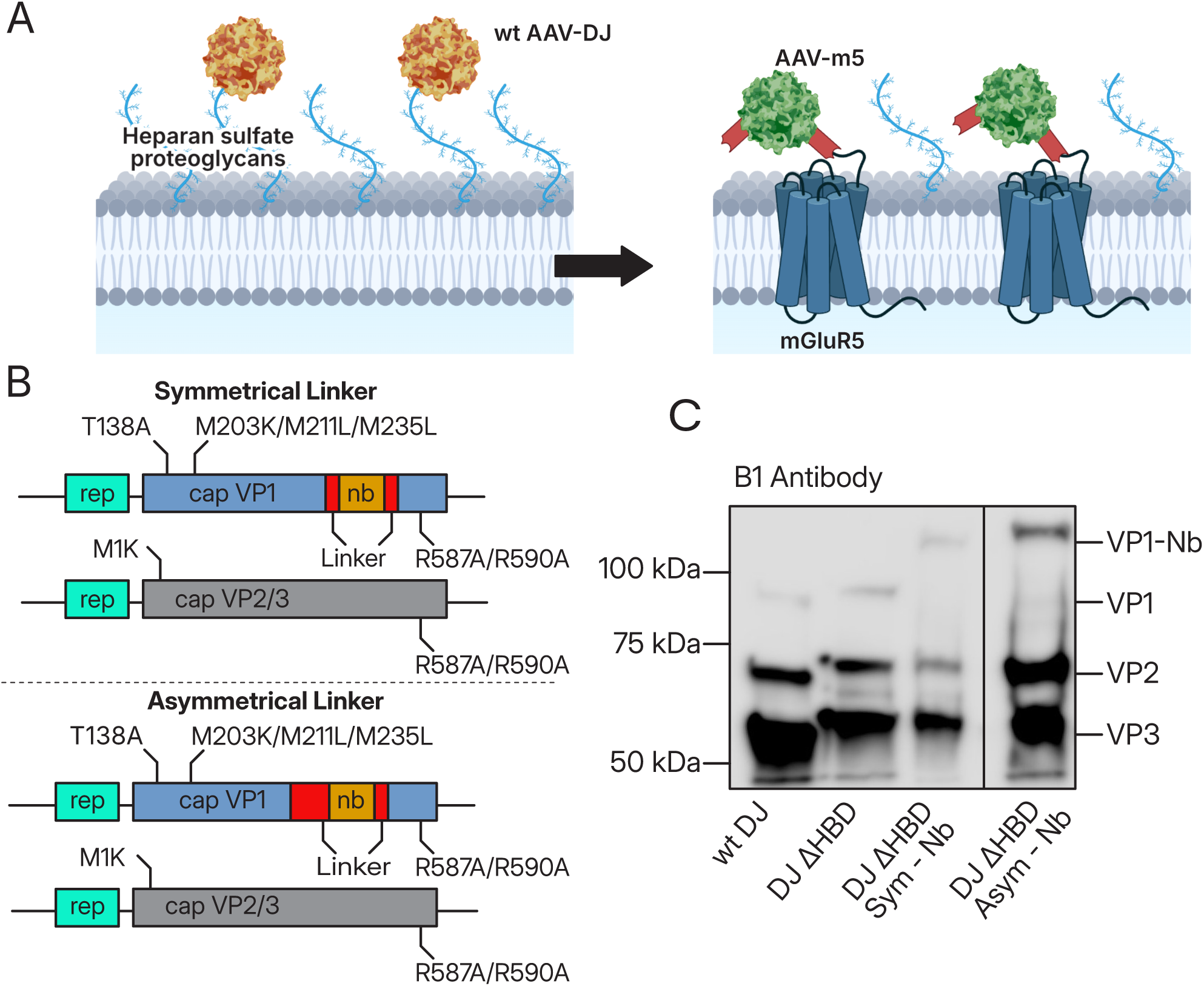
Design and engineering of AAV-m5. **(A)** Schematic of proposed mechanism of re-directed tropism for AAV-m5, replacing heparan sulfate interaction with cell specific mGluR5 interaction. **(B)** Design of plasmids for nanobody incorporation in VP1 of AAV-DJ. **(C)** Immunoblot showing incorporation of nanobody in VP1 of AAV-m5.

### AAV-m5 Rescues Infectivity in an HBD-Null Background

We next evaluated whether the tethered nanobody could restore infectivity to the HBD-null capsid in primary hippocampal neuron cultures. Prior studies have shown that mGluR5 form a protein complex in mouse hippocampus that is also present in hippocampal primary neuronal cultures ^35^. Neurons were transduced at 1x10^4^ vg/cell with AAV variants packaging a GFP reporter, and fluorescence was assessed at seven days post-infection (**Figure 2**). As expected, wild-type virus achieved robust transduction (>99% GFP^+^ cells), while infectivity was abolished in the HBD-null variant. In contrast, incorporation of the mGluR5-targeting nanobody rescued infectivity, even in the absence of heparin binding (48.8 ± 4.21% and 27 ± 4.92% GFP+ cells, for AAV with asymmetric and symmetric nanobody linkers, respectively). Both the symmetrical and asymmetrical linker variants demonstrated comparable transduction efficiencies (p > 0.05, one-way ANOVA with Tukey’s post hoc), indicating that the Nb insertion site tolerates variation in linker geometry without loss of function. To verify the receptor specificity of this rescued tropism, we performed immunofluorescence counterstaining for glial (GFAP) and neuronal (mGluR1/5) markers. Note that the antibodies used to detect mGluR5 show crossreactivity with mGluR1. For AAV-DJ, 30.8 ± 2.81% of cells were negative for mGluR1/5 expression (presumably glia) but showed robust GFP expression. In contrast, AAV-m5-mediated GFP signal colocalized selectively with mGluR1/5-positive neurons (75 ± 7.15% of cells were positive for both GFP and mGluR1/5, while only 0.82 ± 1.18% of mGluR1/5-negative cells showed GFP expression (**Figure 3**)). Taken together, these data indicate that the engineered vector utilizes the targeted receptor for cellular entry.

**Figure 2.**
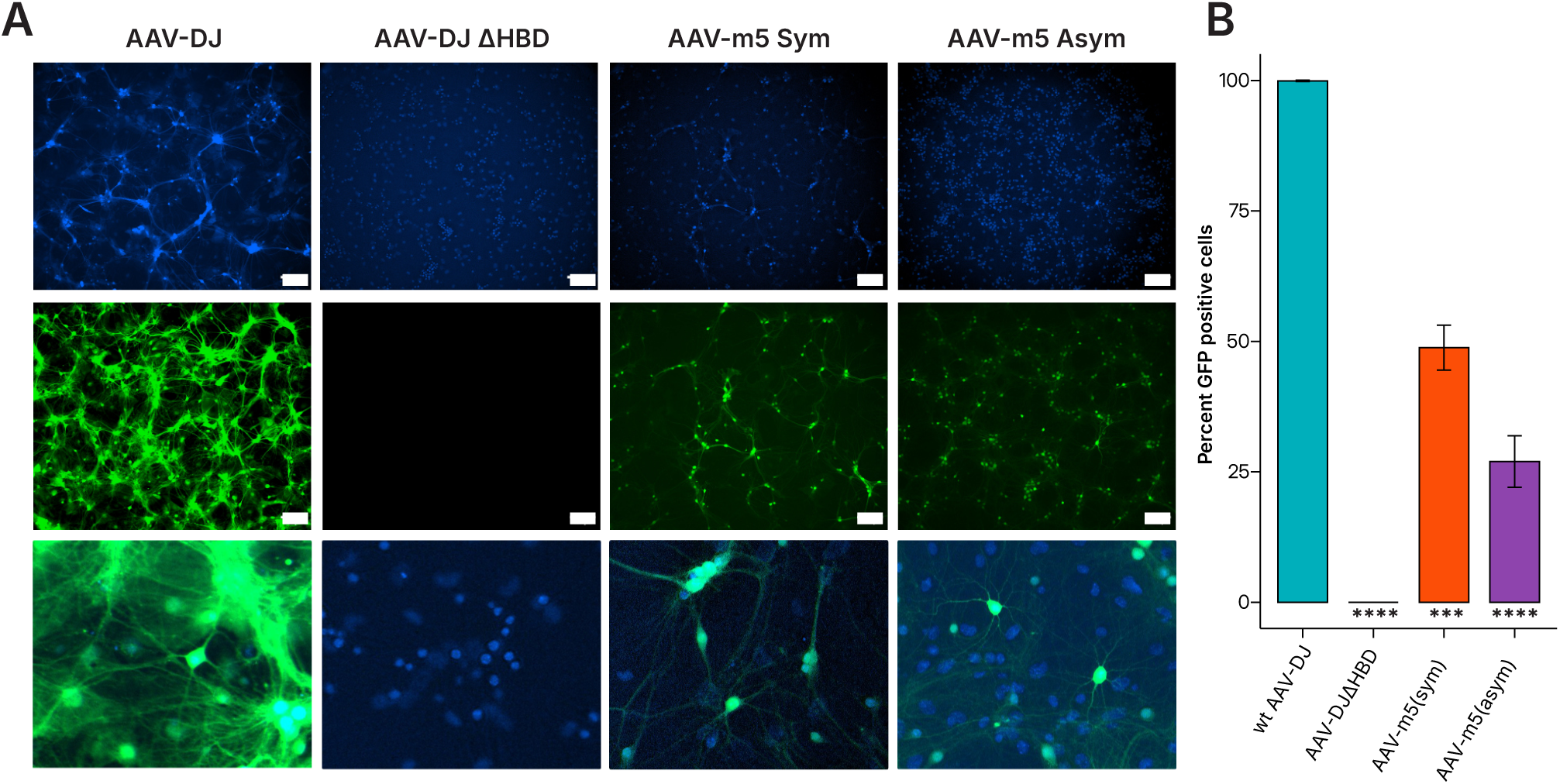
Comparison of AAV-DJ and AAV-m5 variants. **(A)** Virus at multiplicity of infection (MOI) of 1x10^4^ vg/cell was used to transduce primary mouse hippocampal neurons at day 7 *in vitro* and cells were imaged 1 week post transduction. GFP (top row) and Hoescht Nuclear stain (middle row); images taken at 10x magnification, scale bar: 150µm. Bottom Row: Merged GFP/Hoescht and magnified to 40x. **(B)** Quantification of virus transduction as reported by GFP expression. Data are means (n=3-4). Global ANOVA test p value 2x10^-12^. Significance of multiple pairwise comparison with wt AAV-DJ as the control group are indicated as ****: p <= 0.0001 and ***: p <= 0.001.

**Figure 3.**
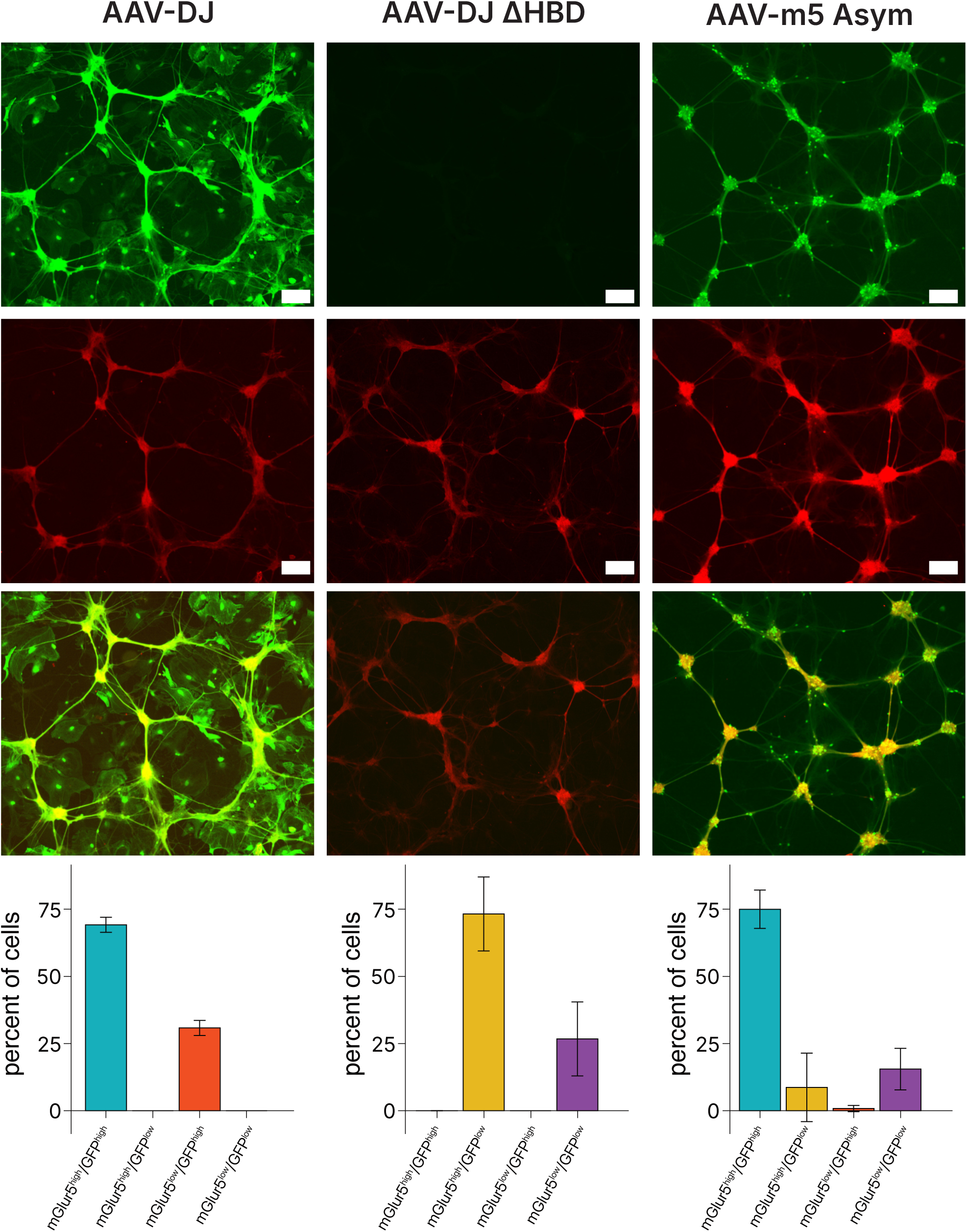
Comparison of AAV-DJ and AAV-m5 variants. Virus at MOI of 1x10^4^ vg/cell was used to transduce primary mouse hippocampal neurons at day 2 *in vitro* and cells were fixed and imaged at day 14 *in vitro*. **(A)** Transgene GFP expression. **(B)** Anti-mGluR1/5. **(C)** Merge. Images taken at 10x magnification, scale Bar: 150µm. **(D)** Quantification of data in **A-C**. N = 5-6. Errorbars indicated standard deviation.

### AAV-m5 Infection is Independent of Heparan Sulfate Binding

To confirm that AAV-m5 utilizes an alternative entry pathway, we performed a heparin competition assay (**Figure 4**). We hypothesized that soluble heparin would saturate the HBD sites of AAV-DJ, thereby blocking interaction with cell-surface HSPGs and reducing infectivity. Consistent with this, AAV-DJ showed a dose-dependent reduction in transduction efficiency across the tested heparin concentration range (0–1111.1 µg/mL; IC_50_ ≈ 37.84 µg/mL). Conversely, AAV-m5 transduction was unaffected across the same concentration range (0.14 ± 1.79% reduction at maximum heparin concentration, p = 0.9008), confirming that its entry mechanism is receptor-dependent and decoupled from canonical HSPG binding.

**Figure 4.**
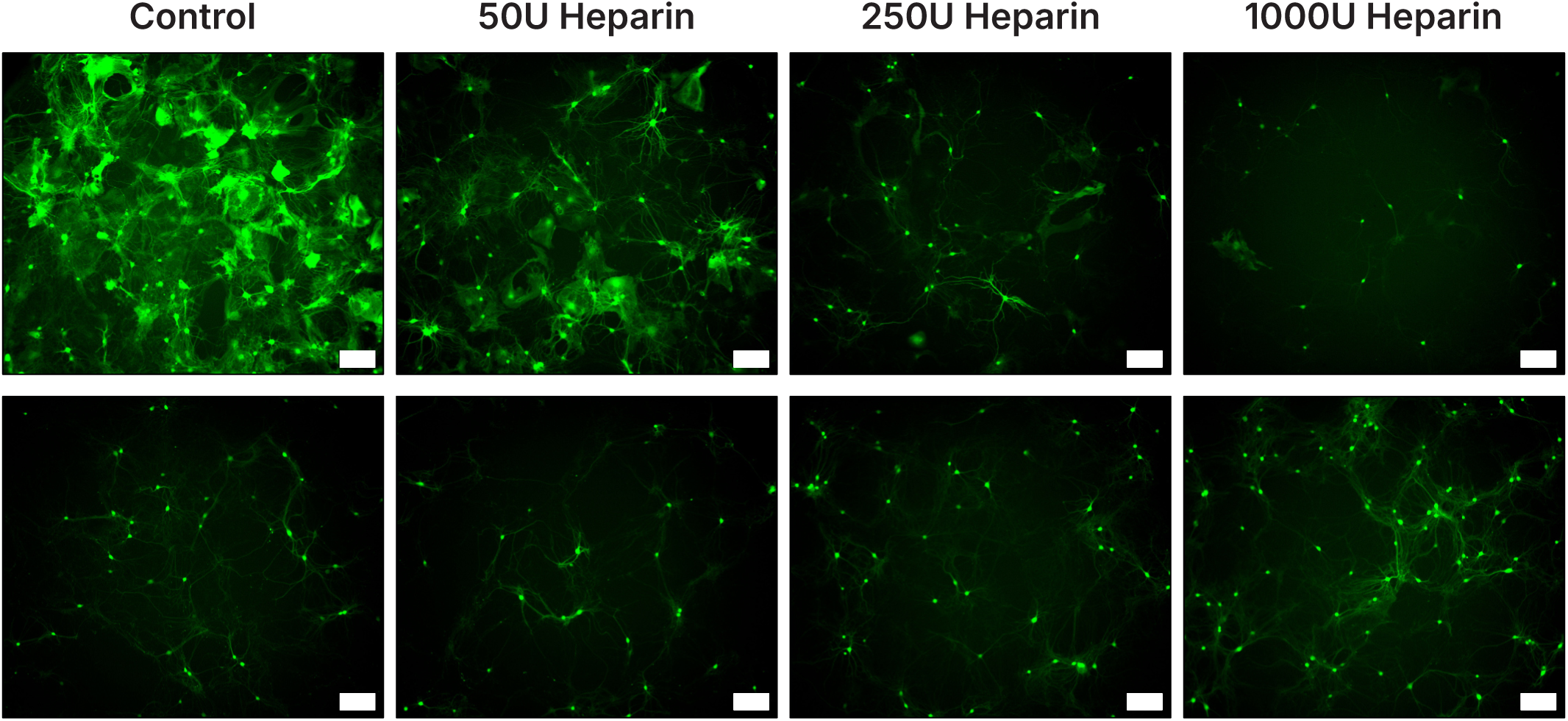
Heparin Competition Assay for wt AAV-DJ and AAV-m5. Virus at MOI of 1x10^4^ vg/cell was incubated with varying heparin concentrations for 30 minutes before transducing primary mouse neurons at day 7 *in vitro*. Media was replaced 2 hours post transduction and cells were imaged 2 weeks post transduction. Top Row: Wild-Type AAV-DJ virus; bottom Row: Nanobody virus (AAV-m5). Images taken at 10x magnification; scale bar: 150µm.

### Targeted Tropism Bypasses the Need for Cell-Specific Promoters

Current strategies for achieving cell-type-specific transgene expression frequently rely on restrictive transcriptional promoters, which can be large, weak, or both ^13–15^. We investigated whether the physical targeting conferred by the mGluR5 nanobody could achieve equivalent specificity in the context of a strong, constitutive promoter (CAG), relative to a neuron-specific (hSyn) or astrocyte-specific (GFAP) promoter (**Figure 5A**). Primary hippocampal neurons were transduced at an MOI of 1x10^5^ vg/cell on DIV7 and fixed 7 days later, on DIV14. Two key findings emerged. First, AAV-m5 packaged with the GFAP promoter yielded negligible GFP expression, confirming that the vector does not efficiently transduce non-neuronal cells despite the activity of the glial promoter in that cell type (**Figure 5B**). Second, the transduction efficiency of AAV-m5 under the constitutive CAG promoter was stronger than that of AAV-DJ driven by the hSyn promoter: 92.1 ± 2.7% vs. 49.8 ± 3.3% of NeuN-positive cells were also GFP-positive (p = 6.03x10^-13^, 2-sample test for equality of proportions with continuity correction). Together, these data suggest that engineered capsid tropism can effectively confine transgene expression to target cell populations, potentially circumventing the packaging size constraints and reduced expression associated with cell-type-specific promoters.

**Figure 5.**
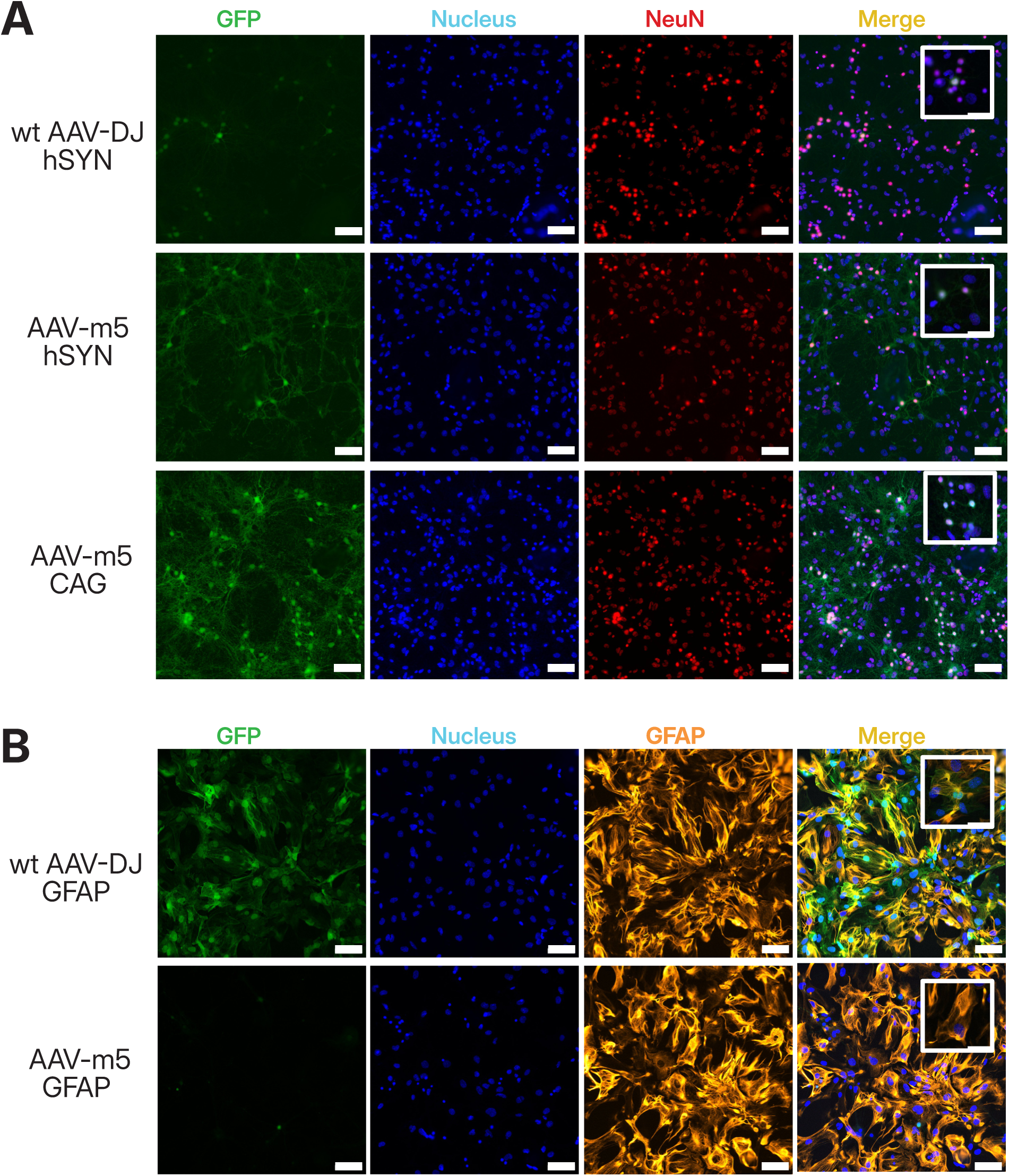
Immunocytochemical analysis of Wild-Type (AAV-DJ) and Nanobody-engineered AAV (AAV-m5). **(A)** Virus packaging a GFP transgene driven by a neuron specific human synapsin (hSYN) promoter was used to transduce primary mouse neurons on day 7 in vitro at a multiplicity of infection of 1x10^5^ vg/cell. Cells were fixed using formaldehyde, stained with neuronal marker (NeuN) and counter stained with the DAPI (Nucleus). **(B)** Virus packaging a GFP transgene driven by a glia specific promoter (GFAP) was used to transduce primary mouse neurons on day 7 in vitro at a multiplicity of infection of 1x10^5^ vg/cell. Cells were fixed using formaldehyde, stained with glial marker (GFAP) and counter stained with DAPI (Nucleus). Scale Bar: 100µm; inset cale bar: 50µm

### Enhanced Performance of AAV-m5 *In Vivo*

Finally, we evaluated the *in vivo* performance of AAV-m5 following stereotactic injection into the mouse dorsal hippocampus, with AAV-DJ delivered into the contralateral hemisphere of the same animals as an internal comparison (1.8 x 10^9^ vg in 250 nL per hemisphere). Because both vectors are therefore represented in every coronal section, this design controls for tissue processing, section thickness, and imaging conditions. Brains were harvested 2 weeks post-injection and coronal cryosections were processed for imaging. At equivalent input titers, AAV-m5 outperformed AAV-DJ, achieving an 8-fold greater proportion of transduced cells within the hippocampus (18.4 ± 1.1% vs. 2.4 ± 0.4%, p = 0.0001764, one-way ANOVA followed by Tukey multiple comparisons of means). These results indicate that nanobody-mediated receptor targeting improves transduction efficiency *in vivo* compared to standard heparin-binding vectors, though the relative contributions of altered cell-surface interactions, diffusion properties, and endosomal processing to this enhancement remain to be determined.

Beyond overall efficiency, the two vectors produced qualitatively distinct spatial distributions. In wild-type animals, AAV-m5 transduction extended across essentially the entire dorsal hippocampal formation (**Figure 6**). GFP signal filled the pyramidal cell layer as a continuous, strongly labeled band running from CA1 at the injection site through CA2 and into CA3, and extended into the molecular layer of the dentate gyrus. Labeling was also prominent in the fasciola cinerea. Outside the hippocampus proper, GFP-positive neurons were present in layer 6 of the adjacent somatosensory cortex, immediately across the corpus callosum from CA1, and in the ventral retrosplenial area. Transduction in the contralateral, AAV-DJ-injected hemisphere was substantially weaker throughout and did not reproduce this laminar organization.

**Figure 6.**
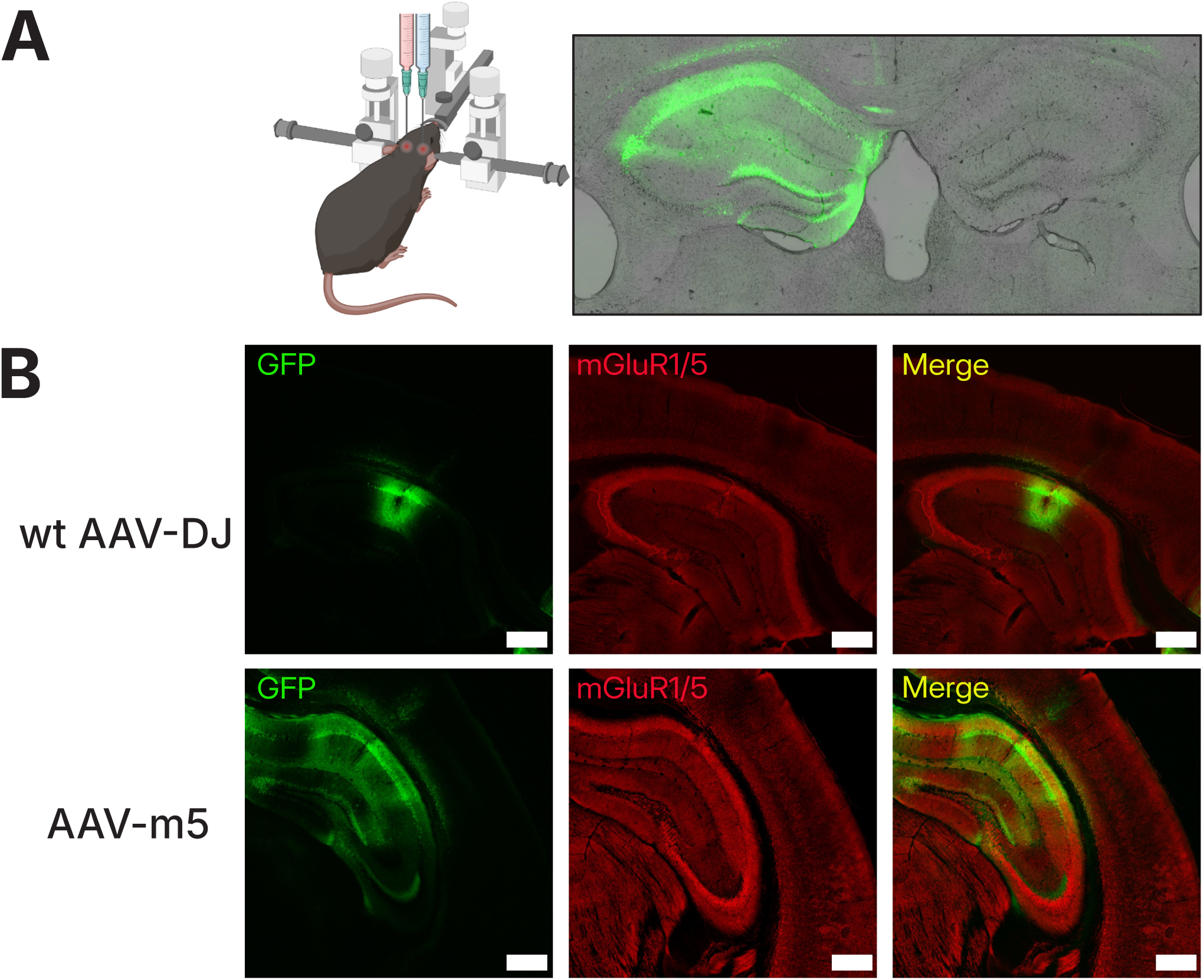
*In vivo* validation and tropism of AAV-m5 in mouse dorsal hippocampus. **(A)** Representative coronal brain sections demonstrating successful virus transduction 14 days post-injection. For comparison, wildtype AAV-DJ was administered to the right hemisphere and AAV-m5 to the left via bilateral stereotactic injection. **(B)** High-magnification immunohistochemistry of fixed hippocampal slices counter stained for mGluR1/5 expression to assess colocalization. Scale bar: 400µm.

To test whether AAV-m5 tropism depends on the intended receptor rather than on a residual capsid-host interaction, we delivered AAV-m5 to the dorsal hippocampus of *Grm5*-null mice at matched dose and coordinates and compared the resulting expression pattern to that obtained in wild-type animals. In *Grm5*-null animals, expression was markedly reduced in both intensity and spatial extent, remaining largely confined to CA1 in the immediate vicinity of the injection tract (**Supplemental Figure 2**). No expression was detected in CA2, CA3, the dentate gyrus molecular layer, the fasciola cinerea, or the retrosplenial cortex. The single extrahippocampal compartment in which labeling persisted was layer 6 of somatosensory cortex directly across the corpus callosum from CA1, where scattered GFP-positive neurons were observed as in wild-type tissue.

## Discussion

The development of precision gene therapy vectors relies on decoupling viral tropism from its natural cell-surface entry mechanisms. Over the past decade, several engineering approaches have emerged to address this challenge, largely focusing on engineerable structural hotspots exposed on the AAV surface ^36–38^. Structural analyses have identified nine variable regions (VR1-9) on the capsid, among which VR4 and VR8 have proven particularly amenable to engineering that redirects viral tropism. Specifically, the T456 residue located within VR4 of the AAV-DJ capsid serves as a structural benchmark for targeted insertions, making it a position for the engineering approaches detailed here ^29, 33, 39, 40^.

Prior modular targeting platforms have relied on the incorporation of small peptide tags into these regions, followed by chemical or enzymatic conjugation of targeting moieties such as DARPins ^41^, bispecific antibody systems ^42, 43^ and SpyTag systems ^44^. However, these multi-component systems require additional post-production steps that pose significant translational challenges regarding manufacturing efficiency, batch fidelity, and in vivo durability.

More recently, the genetic display of nanobodies has emerged as an elegant, single-step alternative. Combining low molecular weight (∼15k Da) with antibody-like binding affinity and minimal immunogenicity, nanobodies can be directly encoded within the AAV cap gene, bypassing the need for post-production assembly completely. While previous work has demonstrated the versatility of this platform to target surface receptors such as CD38, CD4 and FAP ^45, 46^, we have extended this technology to CNS-relevant G-protein coupled receptors (GPCR). By successfully targeting mGluR5 *in vivo*, this work establishes a robust framework for overcoming historical difficulties of cell-type specific delivery within the highly complex architecture of the brain.

### Ablating Endogenous Tropism as a Prerequisite for Precision Targeting

In our view, silencing the vector’s native attachment mechanism was the necessary first step toward precision rather than refined targeting. AAV-DJ is a shuffled chimera derived from a library of eight parental serotypes and is most closely related to AAV2, AAV8 and AAV9; it exhibits excellent transduction kinetics both *in vitro* and *in vivo* ^5^. Although AAV-DJ differs from AAV2 at roughly sixty capsid positions, it retains the canonical heparin-binding domain (HBD) inherited from that lineage, and with it AAV2’s dependence on cell-surface heparan sulfate proteoglycans (HSPGs) as a primary attachment co-receptor ^5, 47^. Systematic charge-to-alanine mutagenesis of the AAV2 capsid localized this property to a cluster of basic residues at the three-fold protrusion, with arginines 585 and 588 (corresponding to R587 and R590 in AAV-DJ numbering) carrying most of the heparin affinity ^48, 49^. Consistent with that mapping, we found that the R587A/R590A double substitution was sufficient to abolish the capsid’s heparin affinity outright, and the resulting HBD-null vector was essentially non-infectious in primary hippocampal cultures, suggesting a clean slate for targeted engineering ^33^.

Mechanistically, there are two main reasons to regard HBD ablation as a prerequisite for precision targeting. The first is competitive. HSPGs are displayed on virtually every cell in the brain and are further embedded throughout the extracellular matrix, at a surface density that exceeds that of any single receptor by orders of magnitude. A targeting moiety grafted onto a capsid that retains high-affinity glycan binding therefore competes against an overwhelming excess of non-specific attachment sites, and the resulting tropism is additive rather than selective. This is why receptor-ablated scaffolds have been a recurring requirement in the DARPin and nanobody retargeting literature, where specificity emerges only once the parental binding activity has been removed ^23, 24, 27^.

The second consideration is pharmacokinetic, and it bears directly on our *in vivo* results. Because HSPGs are distributed throughout the parenchyma and the extracellular matrix, they act as a diffuse sink that immobilizes heparin-binding particles near the injection site. Both pharmacological competition with soluble heparin and genetic ablation of the heparin-binding residues expand the volume of distribution and the extent of AAV2-mediated gene transfer in rodent and non-human primate brain ^50–55^. Our HBD-null scaffold therefore inherits a distribution advantage that is independent and additive to the specificity conferred by the displayed nanobody. We will return to this point when we discuss the spatial footprint of AAV-m5 *in vivo*.

Consistent with these requirements, introducing the anti-mGluR5 nanobody at T456 rescued transduction and confined viral entry almost exclusively to receptor-positive cells, and heparin competition confirmed that AAV-m5 entry is decoupled from background HSPG interaction rather than merely supplemented by a second binding event.

One asymmetry deserves comment. In the studies cited above, heparan sulfate-ablated AAV2 retained transduction *in vivo*, whereas our HBD-null AAV-DJ was essentially non-infectious in dissociated hippocampal culture. We do not think these observations are in conflict. The structure and organization of the extracellular matrix in dissociated cultures is severely altered compared to intact brain tissues, so the sink effect that penalizes heparin-binding capsids *in vivo* is largely absent. Under those conditions, glycan attachment is the dominant route to the cell surface and removing this interaction is correspondingly costly with respect to transduction efficiency. AAV-DJ also binds heparin with higher affinity than AAV2, so an equivalent mutation may impose a larger relative penalty. The practical consequence is that the *in vitro* HBD-null phenotype is mostly a test of receptor-mediated rescue, not a prediction of *in vivo* behavior.

Finally, ablating glycan attachment does not remove the downstream machinery on which AAV entry depends. The binding site for the universal AAV receptor AAVR (KIAA0319L), which contacts the side of the three-fold protrusion, is structurally unperturbed in AAV-m5 ^6, 56^, as are the surfaces implicated in engagement of secondary factors such as FGFR1 ^57^. Residual AAVR dependence is therefore almost certainly present, and it remains to be established whether AAV-m5 uses the target receptor for both attachment and internalization or whether AAVR engagement is still required for endosomal trafficking and nuclear delivery. Given AAVR’s role in post-entry sorting, the interplay between engineered GPCR attachment and native AAVR trafficking is a natural direction for mechanistic follow-up studies that test AAV-m5 in AAVR-null neurons.

### mGluR5 as a CNS Targeting Handle: Receptor Biology and Therapeutic Rationale

mGluR5, encoded by GRM5, is a compelling molecular handle for targeted gene delivery to the CNS on three independent grounds: the breadth and neuronal restriction of its expression, the accessibility of its extracellular domain, and as a therapeutic target itself.

Expression is broad among neurons and restricted to them. mGluR5 is expressed across the principal cell populations of the forebrain, including pyramidal neurons throughout the hippocampal CA fields, dentate granule cells, cortical pyramidal neurons and striatal medium spiny neurons, with immunoreactivity concentrated in somatodendritic compartments ^58, 59^. This breadth of expression is advantageous for promoter-independent neuronal targeting, as AAV-m5 can stand as a capsid-level substitute for pan-neuronal promoters such as hSyn for most forebrain neurons.

An additional aspect of mGluR5-targeted transduction as a substitute for a neuron-specific promoter is that its expression is effectively confined to neurons in the mature brain. Astrocytic mGluR5 is developmentally regulated: it becomes undetectable after the third postnatal week in mouse and was not detected in human cortical astrocytes ^60^. The receptor therefore supplies, at the level of physical entry, the same neuron-versus-glia discrimination that hSyn supplies at the level of transcription and our promoter-comparison data directly support this conclusion. The primary cultures used here derive from neonatal tissue and therefore fall within the developmental window in which astrocytes still express mGluR5. Glial transduction by AAV-m5 was nonetheless negligible, so the neuron-versus-glia discrimination observed *in vitro* cannot be attributed simply to the absence of receptor from astrocytes. Two observations instead indicate that transduction reports on receptor density rather than on receptor presence alone. First, GFP expression tracked receptor immunoreactivity cell by cell: AAV-m5 transduced mGluR1/5-positive cells almost exclusively, whereas AAV-DJ transduced a substantial fraction of receptor-negative cells. Second, astrocytes expressing mGluR5 at neonatal levels were nonetheless largely refractory to AAV-m5. This is the behavior expected of a multivalent capsid, in which several displayed nanobody copies must engage receptor simultaneously to achieve stable attachment, and it predicts that transduction efficiency scales non-linearly with surface receptor density. It is noteworthy that astrocytic mGluR5 re-emerges in rodent models of temporal lobe epilepsy and in resected human epileptic tissue ^61, 62^, so tropism may shift in a reactive or epileptic brain.

The mGluR5 extracellular domain is a validated *in vivo* antibody target. The strongest precedent for the strategy pursued here comes from clinical neuroimmunology rather than vector engineering. mGluR5 was identified as the autoantigen in Ophelia syndrome, a reversible limbic encephalopathy associated with Hodgkin lymphoma; patient antibodies recognize mGluR5 in transfected cells and in cultured rat hippocampal neurons ^63^. Passive transfer of patient IgG into mice alters hippocampal mGluR5 levels and produces behavioral changes that reverse on antibody clearance ^64^. Two conclusions follow, each directly relevant to capsid display. First, the mGluR5 ectodomain is accessible to a binding scaffold on living neurons and in intact brain, and that accessibility is sufficient for productive engagement *in vivo*, which is the requirement for a nanobody presented on a viral particle. Second, sustained occupancy of this ectodomain is functionally consequential, which reinforces the need for the long-term receptor-occupancy studies outlined below.

Furthermore, mGluR5 is a core regulator of synaptic plasticity. mGluR5-null mice show impaired learning and reduced CA1 long-term potentiation ^65^. As such it is among the most intensively pursued CNS drug targets of the past two decades, with programs spanning fragile X syndrome, addiction, levodopa-induced dyskinesia, epilepsy, PTSD and major depressive disorder. AAV-m5 therefore delivers cargo to a neuronal population that is already the focus of sustained drug-development effort and does so through the same receptor those programs target. That convergence cuts both ways: it raises the possibility of combining delivery with modulation, and it makes pharmacological silence of the displayed nanobody a design requirement rather than an incidental property.

The structural rationale for targeting mGluR5 is strong, as well. This receptor is an obligate homodimeric family C GPCR whose large bilobed Venus flytrap domain is linked through a cysteine-rich domain to the seven-transmembrane bundle, projecting the ligand-binding module well above the plane of the membrane ^34^. This architecture is favorable for capsid display in a way that short extracellular loops are not: the epitope sits atop an extended stalk, sterically accessible to a nanobody presented on a 25 nm particle, and the obligate dimer offers the possibility of avidity from adjacent nanobody copies on the capsid surface. mGluR5 is further concentrated in a perisynaptic annulus at the margin of the postsynaptic density at asymmetric excitatory synapses, with a decreasing extrasynaptic gradient ^66, 67^, placing a substantial receptor pool outside the synaptic cleft and within reach of a diffusing particle.

Another beneficial property of this mGluR5 target is its propensity for agonist-driven internalization ^68–70^, which may shift vector dynamics from passive receptor-mediated attachment toward active endocytic uptake. While internalization could transiently reduce infectivity by depleting surface receptor, mGluR5 recycles within hours ^71, 72^. Whether nanobody binding itself drives internalization, and whether it perturbs the recycling cycle, remain open questions. They deserve direct measurement because they bear on transduction kinetics and because chronic receptor depletion would be an adverse effect in its own right.

Selectivity against mGluR1. The Nb43 clone used here was raised and structurally characterized against the mGluR5 extracellular domain, where it binds a surface formed by Helix L and the L-M loop and behaves as a positive allosteric modulator ^34^. Alignment of the ten reported contact positions shows only four conserved between mGluR5 and mGluR1, with three proline substitutions across the segment. These changes are likely to alter loop backbone conformation independently of side-chain identity. Our *in vivo* data support the functional consequence of that divergence. Following delivery to Grm5-null dorsal hippocampus, transduction was confined to CA1 near the injection tract despite robust labeling of the dentate gyrus molecular layer in wild-type tissue, with no detectable expression in the dentate gyrus - the subfield in which mGluR1 transcript is most abundant and mGluR5 least ^59, 73^. Because GRM1 splice variants diverge only in their intracellular C-terminal tails and present an identical extracellular domain, this absence cannot be ascribed to isoform-specific loss of the epitope. That the two ectodomains are immunologically distinguishable is independently established: sera from patients with mGluR5 encephalitis and from patients with mGluR1-associated cerebellar ataxia discriminate between the two receptors in transfected cells ^63^. We therefore describe AAV-m5 as an mGluR5-directed vector and regard the residual perisomatic CA1 and adjacent layer 6 signal as most consistent with receptor-independent entry at high local particle concentration, although low-affinity mGluR1 engagement cannot be formally excluded.

Two implications follow. First, that a capsid-displayed nanobody discriminates between paralogues sharing a substantial fraction of contact residues sets a useful lower bound on the resolution achievable by this approach; subtype selectivity within a closely homologous receptor family is a more demanding requirement than pan-family binding, and one that peptide-insertion and glycan-based targeting strategies are poorly positioned to meet. Second, potentially multiple copies of a positive allosteric modulator displayed on a capsid surface constitute a pharmacological agent as well as a targeting ligand, and local potentiation of mGluR5 signaling at the injection site should be evaluated. Because the nanobody is genetically encoded, signaling-silent clones can be substituted without any change to vector architecture, making such refinement a matter of screening rather than redesign.

### Capsid-Level Targeting as a Functional Substitute for Restrictive Promoters

By conferring precise physical tropism, the AAV-m5 platform offers an alternative to the conventional paradigm of transcriptional gatekeeping. Comparing the highly active constitutive CAG promoter against restricted cell-type-specific promoters (hSyn and GFAP) provides proof of concept that capsid-level engineering can functionally replace promoter-level restriction, and this substitution yields three practical advantages.

The first is payload capacity. The AAV genome imposes a hard ceiling of approximately 4.7 kb, and the constraint is sharper than the nominal figure suggests: systematic analysis places the effective packaging window at 4.1–4.9 kb, with efficiency falling steeply beyond 5.2 kb; oversized genomes are packaged as heterogeneous 5’-truncated species ^74, 75^. Regulatory elements consume this budget directly: the CAG cassette occupies roughly 1.7 kb, hSyn approximately 0.5 kb, the truncated gfaABC1D element approximately 0.7 kb, and full-length GFAP regulatory regions upwards of 2 kb. Moving specificity from the genome to the capsid returns that space to the therapeutic transgene, which matters most for exactly the cargoes that are hardest to accommodate, including base and prime editors, optogenetic reagents, or multi-element regulatory circuits.

The second advantage is expression strength. Cell-type-specific regulatory elements typically drive weaker transcription than constitutive promoters, forcing a trade-off between specificity and transgene level ^76–78^. Our data illustrate the size of that penalty in a single experiment: AAV-m5 under CAG transduced a substantially larger fraction of NeuN-positive neurons than wild-type AAV-DJ under hSyn, while producing negligible expression under GFAP. The capsid-targeted vector did not trade efficiency for specificity, instead it improved both simultaneously.

The third advantage is orthogonality, and it is the one most relevant to therapeutic design. Capsid targeting and transcriptional restriction fail in different ways. A promoter fails when it is transcriptionally leaky in a non-target cell, a behavior that depends on cell state, developmental stage and vector dose. A targeted capsid fails when an off-target cell expresses the receptor or when residual low-efficiency entry occurs by a receptor-independent route. Because the two failure modes are largely independent, layering them should synergize rather than merely add specificity. Furthermore, a third orthogonal layer, such as miRNA-based de-targeting through 3’UTR target sites, can be stacked on top at negligible cost in packaging capacity. For applications with a low tolerance for off-target expression, a dual- or triple-layered architecture combining the engineered capsid with a minimal cell-specific promoter may therefore prove the most stringent cell-type specific targeting strategy.

A final point concerns translatability. Directed evolution of capsids remains a powerful route to novel CNS tropism, but their selected phenotypes can depend on host factors that do not generalize: AAV-PHP.B and AAV-PHP.eB cross the blood-brain barrier through the GPI-anchored protein LY6A, and their striking CNS transduction is restricted to the mouse strain in which they were selected ^8, 17, 79^. Rational nanobody display inverts this dependency. The targeting determinant is a defined, conserved, human-expressed receptor chosen a priori, and the binding interaction is characterized independently of the vector. Species differences in receptor sequence become a testable property of a single 15 kDa module rather than an emergent property of the whole capsid.

### In vivo Performance: Enhanced Efficiency and Spatial Spread

Delivered to dorsal hippocampus, AAV-m5 produced markedly higher neuronal transduction than wild-type AAV-DJ, together with expression extending across the full dorsal hippocampal formation rather than remaining concentrated at the injection site. We considered two explanations for this expanded footprint.

The first invokes enhanced axonal transport, a phenomenon well documented for neurotropic viruses and proposed as the driver of retrograde propagation by rAAV2-retro ^18^. The second, and more parsimonious, explanation is escape from the HSPG sink: by ablating the heparin-binding domain we remove the interaction that immobilizes conventional AAV2-lineage particles in the extracellular matrix near the injection site, permitting broader distribution before productive engagement of a target cell. There is direct precedent for this mechanism: pharmacological competition with soluble heparin and genetic ablation of the same residues expand the distribution of AAV2 in rodent and non-human primate brain ^50–52, 55^.

Three features of our *in vivo* comparison support an mGluR5-dependent entry mechanism. First, the wild-type distribution recapitulates the reported anatomical distribution of the receptor rather than the diffuse parenchymal spread expected of a heparan sulfate-binding vector: labeling followed principal cell layers and their dendritic fields across the CA subfields, the dentate gyrus, the fasciola cinerea and retrosplenial cortex, all of which express mGluR5, with immunoreactivity concentrated in somatodendritic compartments ^58, 59^. Second, genetic deletion of the receptor reduced both the magnitude and the footprint of transduction, establishing that the breadth of the wild-type pattern is receptor-dependent rather than a property of the injection or of vector diffusion. Third, the two compartments in which signal persisted in *Grm5*-null tissue – perisomatic CA1 and the immediately adjacent layer 6 – are precisely those exposed to the highest local particle concentration along and around the injection tract, which is the distribution expected of a low-level, receptor-independent entry (e.g., through micropinocytosis) rather than of engagement with an alternative receptor.

### Limitations and Future Directions

Despite the promising performance of the AAV-m5 vector, certain translational limitations remain to be systematically addressed. Our current two-week *in vivo* endpoint provides an accurate snapshot of acute transduction and initial tropism; however, it does not account for the long-term stability of transgene expression, potential host immune responses mounted against the non-human nanobody domain. Long-term longitudinal studies will be vital to assess these safety parameters.

Additionally, while small-scale iodixanol gradient purification configurations successfully yielded highly functional AAV-m5 preparations, the modified capsids exhibited lower viral titers than their wild-type counterparts. Whether this reduction stems from altered packaging dynamics or purification recovery anomalies remains unknown, and evaluating full manufacturing scale-up parity is essential for future development.

Nevertheless, the inherent modularity of this display platform establishes the well-characterized T456 site as a powerful framework for rational vector design. Because the nanobody sequence is genetically integrated, this system operates as a “plug-and-play” architecture. Because mGluR5 defines a broad neuronal address, narrower cell-type restriction awaits binders against surface antigens with correspondingly narrower distributions: the anti-mGluR5 sequence can be swapped for alternative single-domain antibodies tailored to distinct CNS targets, enabling researchers to selectively target cell-type-specific surface antigens unique to interneuron subpopulations, microglia, astrocytes, oligodendrocytes, or even pathological extracellular protein aggregates. Ultimately, validating this modular platform across systemic administration routes, large-animal models, and human *ex vivo* organotypic tissue slices will mark the next major milestones toward clinical realization.

## Methods

### Molecular cloning

All oligos and gene fragments (gBlocks) were obtained from IDT. PCRs were performed using the PrimeSTAR Max DNA polymerase from Takara Bio. PCR products were gel-purified using the Zymoclean Gel DNA Extraction Kit from Zymo Research after analysis on a 1% TAE agarose gel. AAV-DJ-VP1 and AAV-DJ-VP2/3 were obtained by mutating the start codons for VP2/3 (T138A, M203K, M211L, M235L) and VP1 (M1K), respectively. This was followed by site directed mutations to ablate heparin binding (R587A and R590A). Plasmids were cloned via either Gibson cloning using NEB Hi-Fi cloning mix or Golden Gate cloning using NEB’s T4 DNA ligase and appropriate Golden Gate enzymes (BsmBI) and transformed into *E.coli* cells before being plated on LB plates containing carbenicillin at a concentration of 100 µg/mL. OneTaq 2x Master Mix from NEB was used to perform diagnostic colony PCR to screen for positive clones. Colonies corresponding to the positive clones were then inoculated in 3 mL LB media supplemented with 100 µg/mL carbenicillin and plasmids were isolated using the Zyppy Plasmid Miniprep Kit from Zymo Research, according to the manufacturer’s instructions. Plasmids were then verified for successful cloning via the Plasmid-EZ sequencing service provided by Azenta Life Sciences. All plasmids used in this study are listed in **Table S1**.

### Tissue Culture

HEK293AAV cells (Cell Biolabs) were cultured in DMEM (Gibco) supplemented with 10% fetal bovine serum (Gibco), 4.5 g/L D-glucose, L-glutamine, 110 mg/L sodium pyruvate, and 100 U/mL penicillin/100 µg/mL streptomycin (Gibco). Cells were kept in a humidified incubator at 37℃ and 5% CO_2_ and passaged every 2-4 days when reaching 70% confluency. Cells were passaged until passage 10 for AAV production.

### Neuron Culture

Timed-pregnancy female Mice (CD1) were procured from Charles River laboratories. On Day 0, brains were harvested from newborn pups (P0) and dissected in dissection media (35 mM D-Glucose, 20 mL Ky/Mg stock (Working concentration: 1mM Kynurenic acid, 10mM MgCl_2_) in 200mL Hank’s Balanced Salt Solution) to obtain hippocampi. Hippocampi were then collected in a 50 mL falcon tube and digested using 3 mL of Digestion Solution (1.6 mg L-Cysteine hydrochloride, 100 U Papain, 5mL Dissection media, pH 7.4) for 6 minutes at 37°C. Hippocampi were then washed 3 times with Inhibitor Solution (50 mg BSA, 50 mg Ovomucoid trypsin inhibitor, 5 mL Dissection media, pH 7.4). Inhibitor Solution (3 mL) was added and incubated at 37°C for 4 minutes. Inhibitor Solution was then removed and washed 3 times with 1 mL of Plating media, this was transferred to a fresh 50 mL Falcon tube and the volume was made up to 2 mL. This was triturated using a 1 mL pipette and allowed to settle down for 1 minute. 1 mL of supernatant was removed and replaced with 1 mL of plating medium (minimum essential media (MEM) supplemented with 5 g/L glucose, 10 mM HEPES, 100 µg/mL transferrin, 2 mM L-glutamine, 25 µg/mL insulin, 10% HI-FBS, 2% B-27 supplement, pH adjusted to 7.4) before triturating again. Cells were then counted in a hemocytometer, and 30,000 cells were plated in 100 µL plating media per well of a 24 well plate that had been previously coated with Matrigel. Cells were incubated at 37 °C for 2 hours to allow cells to adhere following which 1 mL of warm plating media was added to each well. Depending on the experiment, 1 mL of AraC media (MEM supplemented with cytosine arabinoside (AraC), 5 g/L glucose, 10 mM HEPES, 100 µg/mL transferrin, 4 mM L-glutamine, 12.5 µg/mL insulin, and 5% (v/v) HI-FBS and pH adjusted to 7.4) was added on Day In Vitro (DIV) 2 to reduce proliferation of non-neuronal cells.

### Iodixanol-Gradient AAV Production

AAV was purified using iodixanol gradients according to previously published protocols ^80^. Briefly, 6 million 293AAV cells were seeded into 15 cm dishes in 25 mL of media. 10 dishes were seeded for each gradient production. 48 hours post-seeding, cells were transfected with 47 µg of plasmid DNA per dish using PEI (MW 25,000, Kyfora Bio). For AAV-m5 production, plasmid DNA was split in a 1:1:1:1 equimolar ratio of (i) an Adeno-helper plasmid, (ii) a cargo payload plasmid, (iii) a plasmid encoding the *rep* and *cap* genes for VP1 with the nanobody incorporated, and (iv) a plasmid encoding the rep and cap genes for the remaining complementing VP2 and VP3 proteins. For non-nanobody AAV production, plasmid DNA was split in a 1:1:1 equimolar ratio of (i) an Adeno-helper plasmid, (ii) a cargo payload plasmid and (iii) a plasmid encoding the rep and cap genes for the VP1/2/3 proteins either without HBD mutations (for wild type) or with HBD mutations (for HBD mutants). All plasmid combinations used for virus production are outlined in **Supplementary Table 1.**

72 hours post transfection, cells from 10 dishes were harvested using a cell lifter, washed with phosphate-buffered saline (1x PBS) once, and resuspended in a Benzonase buffer containing 2 mM MgCl_2_, 0.15 M NaCl, and 50 mM Tris-HCl at pH 8.5. Five freeze thaw cycles were then performed on the samples, alternating between 2 minutes in liquid nitrogen and 10 minutes in a water bath at 37°C. This lysed the cells and released AAV particles. Free plasmid and genomic DNA from cells was then digested with benzonase nuclease (Sigma-Aldrich) for 1h at 37°C. Samples were then spun down twice at 4000xg, 4°C to separate cell debris from AAV lysate. This AAV lysate was then loaded into ultracentrifugation tubes and iodixanol (Progen) amounts of 15%, 25%, 40% and 60% were layered beneath it. Tubes were then sealed and subjected to density gradient centrifugation in a Ti 70.1 (Beckman) for 2 hours at 176,400 x g (50,000 rpm) and 4°C. The 40% phase containing AAV particles was then collected post-centrifugation.

### Quality Control of AAV

Iodixanol gradient purified AAV was then assessed for production titers and VP incorporation via quantitative polymerase chain reaction (qPCR) and Western blot, respectively. qPCR was performed as follows: 2 µL of purified AAV was mixed with 42 µL of PBS, 5 µL of Proteinase K Buffer (100 mM Tris-HCl, pH 8.0, 10 mM EDTA, 10% SDS) and 1 µL of Proteinase K (20 mg/mL; Zymo Research). This was then incubated for 30 minutes at 50°C to digest the proteinaceous AAV capsid and release the single-strand DNA (ssDNA) payload. Heat inactivation of Proteinase K was performed at 95°C for 10 minutes. ssDNA was then purified using a DNA Clean & Concentrator-5 Kit (Zymo Research) according to the manufacturer’s instructions. Samples were diluted 1:1000 in nuclease free water and used for qPCR using the PowerUp SYBR Green Master Mix (Applied Biosystems) on QuantStudio 5 Real Time PCR System (Applied Biosystems). Primers specific to the CMV enhancer (forward primer, AAC GCC AAT AGG GAC TTT CC; reverse primer, GGG CGT ACT TGG CAT ATG AT), the GFAP promoter (forward primer, TGG CAG CAT TGG GCT GG; reverse primer, GTG CGG CGG GAA GCA G) or the hSyn promoter (forward primer, CTG ACG ACC GAC CCC GAC; reverse primer, TGA AGC TGG CAG TGC GC) on the ssDNA payload were used together with a plasmid standard at a known concentration to calculate the viral titer in viral genomes per milliliter.

For Western Blots, a known concentration of purified AAV (>5x10^9^ vg/mL) was mixed with 12.5 µL of 4x Laemmli Sample Buffer (Bio-Rad), supplemented with 2-mercaptoethanol. The volume was made up to 50 µL with PBS and denatured at 95°C for 10 minutes. Samples were then spun down and cooled on ice before being loaded and separated on a 7.5% precast polyacrylamide gel (Bio-Rad). 10 µL of Precision Plus Protein Dual Color Standard (Bio-Rad) was loaded as a molecular weight ladder. The gel was run at 120V until the 10 kDa band reached the bottom in Tris/glycine/SDS Electrophoresis Buffer (Bio-Rad). Subsequently, proteins were transferred onto a nitrocellulose membrane (pore size = 0.45 µm, Thermo Scientific) in an ice-cold blotting buffer (25 mM Trizma base, 96 mM glycine, and 20% methanol) for 80 minutes at 110V. Post transfer, the membrane was washed once for 5 minutes with 1x TBS-T buffer (20 mM Tris base, 137 mM NaCl, pH 7.6, 0.05% Tween-20) and then incubated in blocking solution (5% skim milk in 1x TBS-T) for 1 hour at room temperature on a rocker. Primary Antibody B1 (Anti VP1/VP2/VP3 mouse monoclonal, Progen Cat# 65158) was used at a 1:250 dilution in blocking solution. 20 mL of this antibody dilution was added to the membrane and incubated overnight at 4°C. The next day, the membrane was washed four times for 5 minutes each with 1x TBS-T. Secondary antibody (Goat anti-mouse IgG-peroxidase antibody, Sigma Aldrich, A2304) was diluted to 1:25,000 in blocking solution and added to the membrane. This was incubated on a rocker at room temperature for 1 hour. Post-incubation, the secondary antibody solution was discarded, and the membrane was washed four times for 5 minutes each with 1x TBS-T. Subsequently, SuperSignal West Dura Extended Duration Substrate kit solution (Thermo Scientific) was added to the membrane and chemiluminescence signal detected using an Amersham Imager 600 (GE Healthcare).

### Viral Transduction of Primary Neuron Culture

Primary hippocampal neurons were transduced at DIV3 or DIV7 to allow for maturation of neurons and expression of mGluR1/5 receptors ^35^. To ensure optimal cell viability during infection, viral transduction was performed in plating media. Briefly, 200 µL of existing media was aspirated from each well and transferred to a sterile 1.5 mL microcentrifuge tube. Iodixanol-purified AAV were then added to the media and homogenized by gentle pipetting. This mix was then returned to its respective wells. Transduction was performed in triplicate for each condition at multiplicities of infection (MOI) of either 1x10^4^ or 1x10^5^ vg/cell. Post transduction, cells were returned to a humidified incubator at 37°C and 5% CO_2_ for up to 2 weeks.

### Heparin Competition Assay

To assess the dependence of AAV-m5 variants on cell-surface heparan sulfate proteoglycans (HSPGs) for viral entry, a competition assay was performed using Heparin Sodium Salt (Sigma-Aldrich). Heparin was reconstituted in deionized water to a stock concentration of 5,000 U/mL. Serial dilutions were prepared by diluting with 1x PBS. AAV particles at a fixed multiplicity of infection (MOI) were combined with varying concentrations of heparin or PBS vehicle control. The mixtures were incubated for 30 minutes at room temperature to allow for saturation of the viral heparin-binding domains. Prior to transduction, 1 mL of culture media was harvested from each well and stored at 4°C. An additional 200 µL of media was collected and added to the virus-heparin mix. This mixture was then added to the side of the wells to prevent physical disruption of the cell monolayer. Following a 2-hour incubation at 37°C, the media was aspirated and discarded. The previously harvested conditioned media was returned to the respective wells. Transduction efficiency was assessed via fluorescence imaging 2 weeks post-infection.

### Immunocytochemistry Staining

Media from wells was discarded by overturning the plate. Cells were washed twice by adding 400 µL 1x PBS to the walls of the wells and then overturning the plate. 500 µL of 2% PFA (5 mL 2x Cytoskeleton Buffer, 1.25 mL 16% PFA, 3.75 mL water) was added to each well and incubated for 10 minutes in the fume hood. This was then aspirated using a pipette and wells were washed thrice with 1x PBS. To permeabilize, 500 µL of freshly prepared 0.1% Triton-X-100 was added to the wells and incubated at room temperature for 10 minutes. Cells were then washed once with 1x PBS following which 500 µL of Blocking Buffer (5% Normal Goat Serum in 0.1% Triton-X-100) was added and incubated at room temperature for 1 hour on a shaker. Primary Antibodies were prepared by diluting in Blocking Buffer accordingly, Rabbit Anti-mGluR1/5 (R&D Systems PPS079) - 1:500, Chicken Anti-GFAP (Novus Biologicals NBP1-05198) - 1:5000, Rabbit Anti-NeuN (ThermoFisher Scientific 702022) - 1:500, Mouse Anti-mGluR1/5 (ThermoFisher Scientific MA5-27691) - 1:250. Blocking Buffer was removed from the plates and replaced with 250 µL of the primary antibody solution. Plates were then wrapped in parafilm and left in the refrigerator overnight. The next day, secondary antibodies were prepared by diluting in Blocking Buffer accordingly, Goat Anti-Rabbit AF555 (ThermoFisher Scientific A-21428) - 1:200, Goat Anti-Chicken AF647 (ThermoFisher Scientific A-21449) - 1:200, Goat Anti-Mouse AF568 (ThermoFisher Scientific A-11004) - 1:200. Primary antibody solution was removed, and wells were washed thrice with 1x PBS. 250 µL of the secondary antibody was added to the wells and incubated at room temperature on a shaker for 1.5 hours. Secondary antibody solution was then removed and 250 µL of DAPI nuclear stain (1:1000) was added to the wells and incubated for 5 minutes. DAPI solution was discarded and cells were washed thrice with 1x PBS. 10 µL of prolong antifade mounting media was added to each well and coverslips were placed on the cells. This was allowed to air dry in the dark overnight before being imaged.

### Immunohistochemistry staining

Post fixing coronal slices were sectioned and stored in 1x PBS in a 24 well plate. PBS was removed and tissue was permeabilized using Permeabilizing solution (1x PBS + 0.3% Tween-20) for 2 hours on an orbital shaker at room temperature. A blocking solution (10% NGS in Permeabilizing solution) was added and incubated for 1 hour on the orbital shaker at room temperature. Primary antibody solutions were diluted in the blocking buffer as follows: rabbit anti-mGluR5 (1:500), Chicken anti-GFAP (1:5000). 500 µL of this was added to each well and incubated overnight on an orbital shaker in the cold room. The next day, wells were washed with permeabilizing buffer thrice for 10 minutes each on an orbital shaker. Secondary antibody solutions were diluted in blocking buffer (1:200) and added to the wells. This was incubated for 1 hour at room temp on an orbital shaker. Afterwards, the wells were washed thrice with permeabilizing buffer for 10 minutes each on an orbital shaker. DAPI solution was diluted 1:1000 in PBS and added to the wells. This was incubated for 5 minutes. Following this, wells were washed thrice with 1x PBS. Slices were then transferred to glass slides and allowed to air dry in the dark for an hour before adding 15 µL of Prolong mounting media. A cover slip was added, and slides were allowed to air dry for a minimum of 24 hours before being sealed.

### In vivo viral transduction

All animal use was reviewed and approved by the Institutional Animal Care and Use Committee of the University of Minnesota. All experiments involving animals were designed, performed, and reported in accordance with institutional regulations and ARRIVE Guidelines (2.0) ^81^. Intracranial surgeries were performed as described ^82^ with minimal modifications. Briefly, C57BL/6J mice (male, age 8-10 weeks) were anesthetized with isoflurane using a low-flow anesthesia system (Somnoflo, Kent Scientific: Torrington, CT), positioned in a stereotaxic frame (Model 1900, David Kopf Instruments; Tujunga, CA), and the skull leveled. Custom-made microinjectors (33 Ga) were lowered into the skull through the burr holes to the dorsal HPC (from bregma: −2.2 mm A/P, ± 1.6 mm M/L, −1.4 mm D/V). Viral vectors (250 nL) were delivered bilaterally at a speed of 100 nL / minute and the microinjectors were left in place for 10 minutes after infusion to decrease viral diffusion through the injection track. Animals were allowed to recover for 2 weeks before viral expression was assessed. For *post hoc* evaluation of intracranial targeting, mouse brains were extracted, fixed in 4% PFA (in PBS) overnight, and cryoprotected in 30% sucrose (in PBS) for 48 hours. Brains were then sectioned coronally at 50 µm on a freezing sliding microtome (Leica SM 2000 R, Leica Biosystems). Slices containing the dorsal hippocampus were collected and injection locations were assessed by evaluation of viral-driven GFP fluorescence using a BZ-X800 epifluorescent microscope (Keyence). Images for each channel were obtained from multiple focus planes and stitched using Keyence BZ-X800 analysis software. Fluorescence images were overlaid with brightfield images.

## SUPPLEMENTARY MATERIALS

**Supplementary Table 1.**
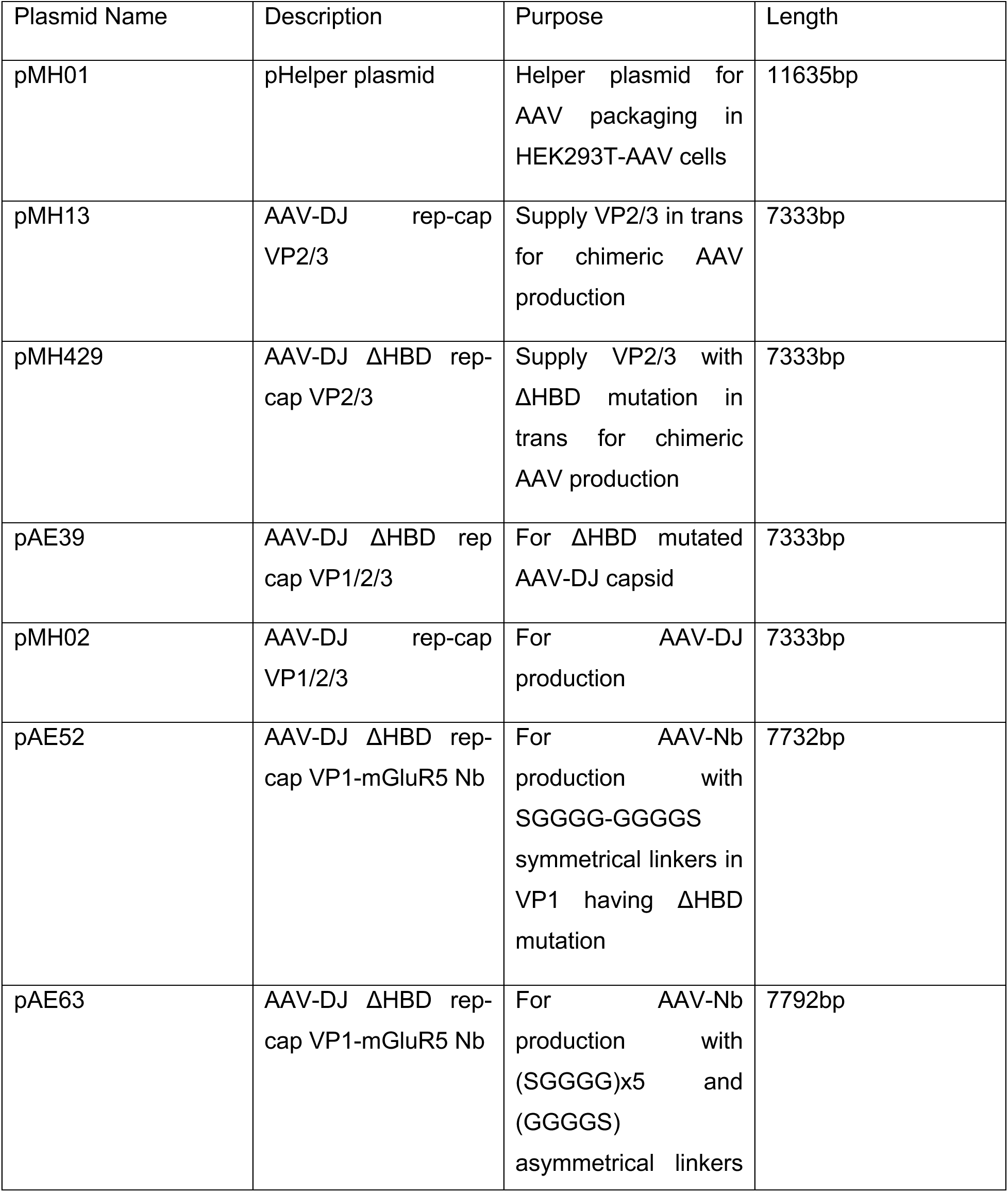
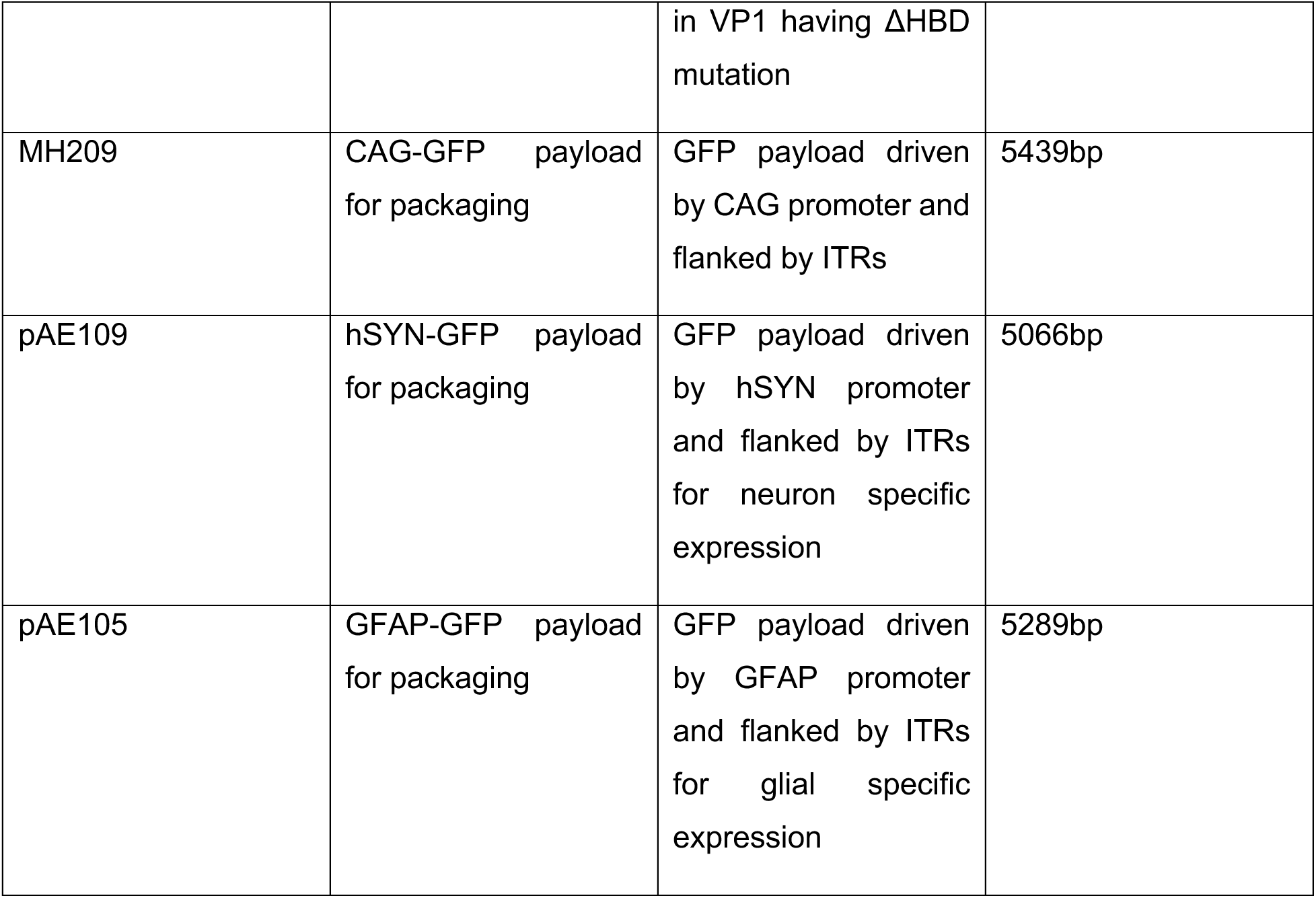
Plasmids used in this study.

**Supplementary Fig 1.**
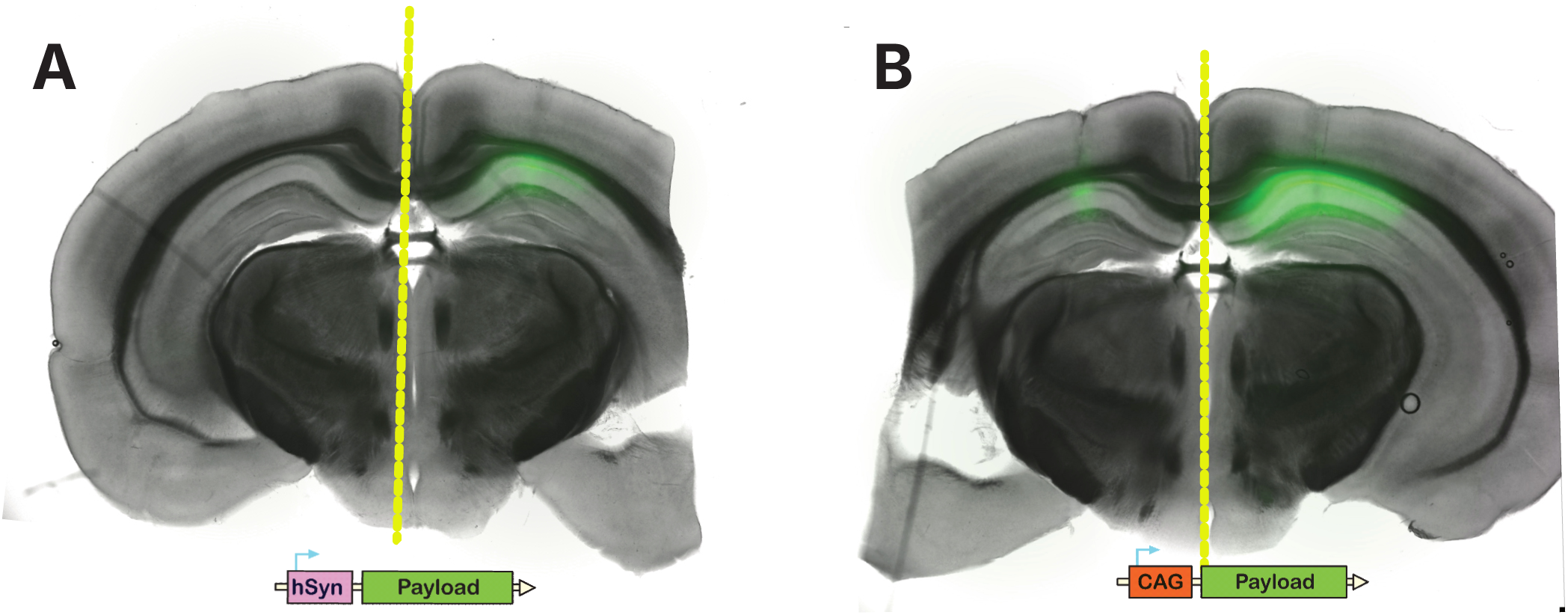
*In vivo* validation of AAV-m5 in mouse hippocampus. **(A)** Comparison of AAV-DJ and AAV-m5 delivering a GFP payload driven by a neuron specific hSyn promoter, Left hemisphere: wt AAV-DJ, Right hemisphere (Notched): AAV-m5. **(B)** Comparison of AAV-DJ and AAV-m5 delivering a GFP payload driven by a constitutive CAG promoter, Left hemisphere (Notched): wt AAV-DJ, Right hemisphere: AAV-m5; 150µm thick coronal slices.

**Supplementary Fig 2.**
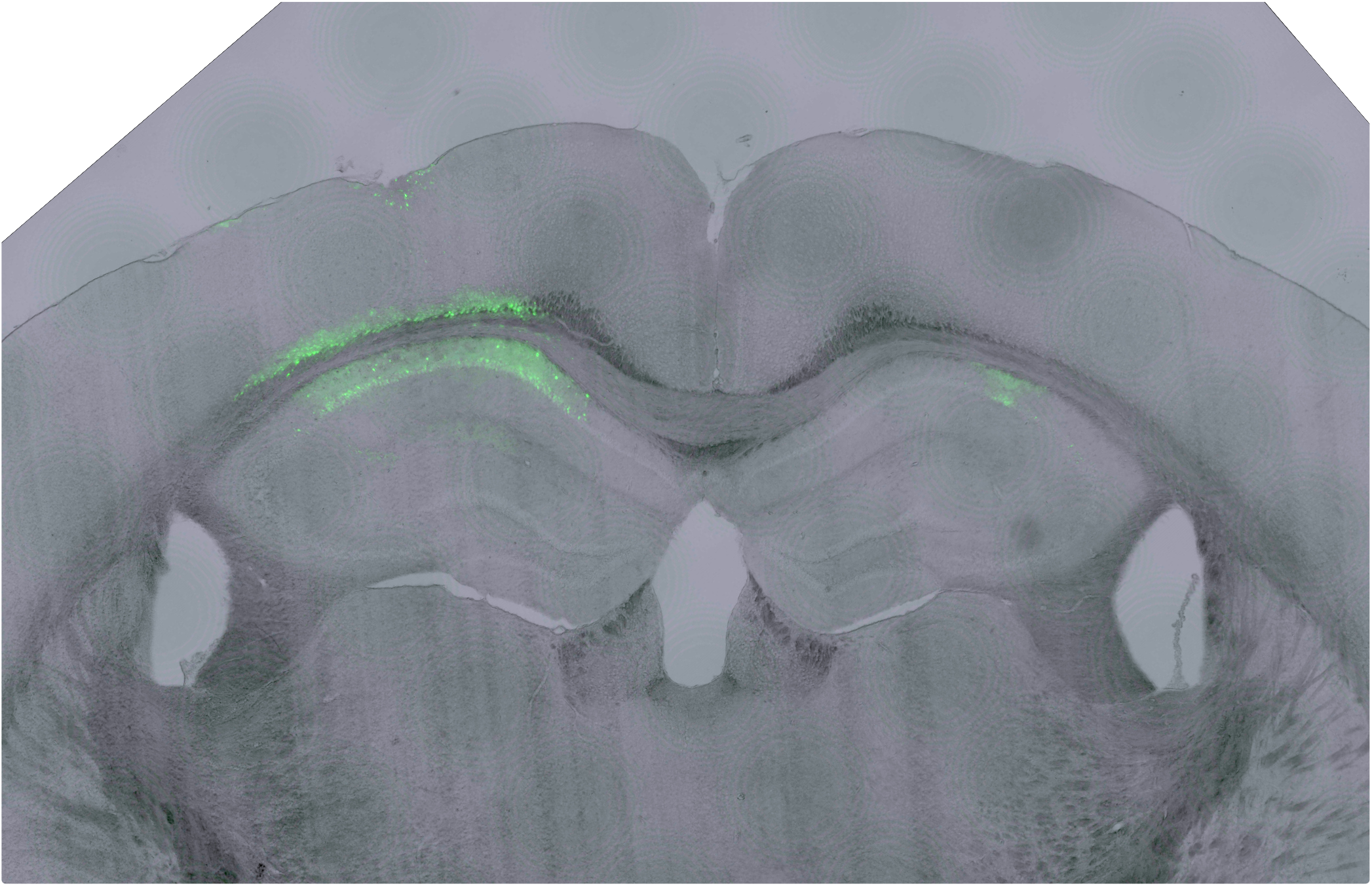
Viral transduction in dorsal hippocampus of GRM5 knockout mice. **(A)** Representative coronal brain section comparing transduction of AAV-m5 (left hemisphere) and wildtype AAV-DJ (right hemisphere) delivering a GFP payload driven by a constitutive CAG promoter.

## Notes

### Competing Interest Statement

The authors have declared no competing interest.

